# Hub taxa are keystone microbes during early succession

**DOI:** 10.1101/2023.03.02.530218

**Authors:** Amanda H. Rawstern, Damian J. Hernandez, Michelle E. Afkhami

**Affiliations:** Department of Biology, University of Miami, Coral Gables, FL, 33146, USA

## Abstract

Microorganisms underpin numerous ecosystem processes and support biodiversity globally, yet we understand surprisingly little about what structures environmental microbiomes themselves. Combining culturing, sequencing, and microbial networks, we identified ‘central’ (highly-connected, hub taxa), ‘intermediate’ (moderately-connected), and ‘peripheral’ (weakly/un-connected) microbes and experimentally evaluated their effects on soil microbiome assembly during early succession. Our results demonstrate central early colonizers significantly (1) enhanced biodiversity, (2) increased recruitment of additional influential, hub taxa, and (3) shaped microbiome assembly trajectories. This work elucidates fundamental principles of network theory in microbial ecology and demonstrates for the first time in nature that central, hub microbes are keystones.

## Main Text

As microorganisms drive global ecosystem processes including decomposition, nutrient cycling, and maintenance of biodiversity^1^, it is imperative that we understand what factors affect how these communities assemble. In addition to abiotic filtering and interactions with macro-organisms^2^, increasing evidence suggests that inter-microbial interactions shape the assembly of microbiomes^3^. However, it remains difficult to identify which microbial taxa play key roles in structuring their communities. Attributes of microbes such as rarity/commonness across a landscape^4^, degree of habitat specialization^5^, and possible role as keystone taxa^6^ have been proposed to be important in understanding which microbes influence community structure. In particular, ongoing debate over the last decade has centered on whether network properties can predict how influential a microbe is within the community. Central taxa (highly-connected “hub taxa” within microbial networks) are theorized to be keystone species that disproportionately structure microbial communities^6^. Current evidence supporting central microbes as keystones is based on computational predictions^7-8^, synthetic communities^9-10^, and laboratory manipulations^11-12^, however no studies have experimentally tested if hub taxa govern microbiome community dynamics in nature. To address this gap we bring together microbiome sequencing, culture collections, network analysis, and field microcosms to identify ‘central’, ‘intermediate’, and ‘peripheral’ microbes within microbiome networks and, for the first time, experimentally evaluate their effects on early microbial community assembly in a native ecosystem.

We isolated and identified with Sanger-sequencing 66 microbial cultures from soils collected across 64 Florida Rosemary Scrub sites^13^. This pyrogenic ecosystem naturally experiences succession following fire sterilization of the top layer of soil, ideal for the study of community assembly during early succession^14^. We used *BLAST* to compare the isolated taxa’s sequences against the NGS microbiome-wide Florida scrub dataset (from 103 sites)^13^ and microbiome network to match the isolated microbes to their ecological attributes of landscape rarity, habitat specialization^15^, and network centrality tier^16^ (attributes described in Fig 1 and Supplementary Methods). The cultured microbes spanned the natural variation in all three ecological attributes (Fig1 A-C), making them excellent representatives of the natural community. We conducted a manipulative field experiment, inoculating 100 sterile soil microcosms with one of 20 isolated, characterized microbes. We then monitored how the ecological attributes of these “early colonizers” affected microbiome assembly in their native habitat. To investigate early succession, 300 soil samples were collected from the field across three time points (1 day, 7 days, and 14 days) and then sequenced to characterize prokaryotic diversity, composition, and recruitment (Supplementary Methods).

**Fig 1.**
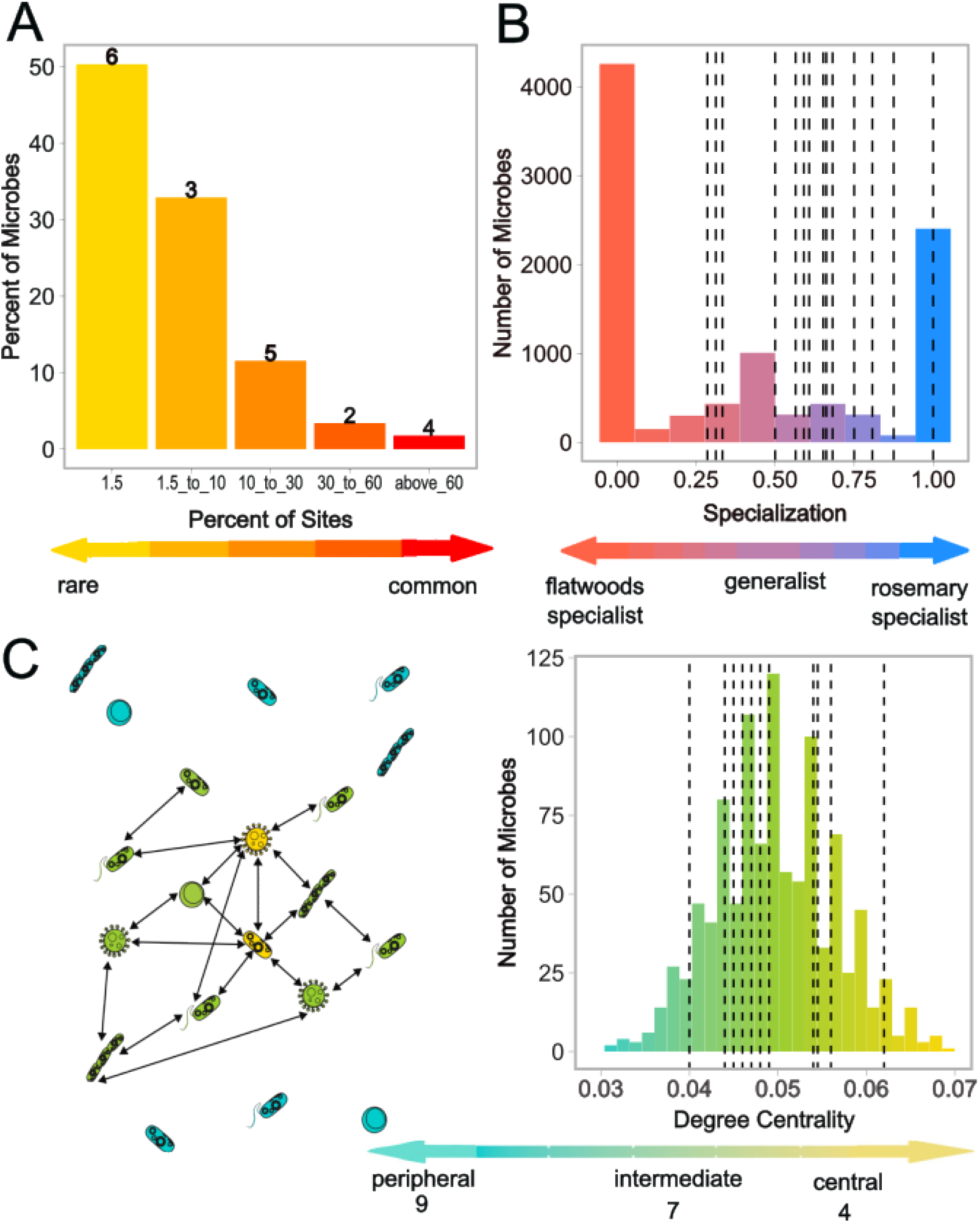
Ecological attributes of isolated microbial taxa used in the field experiment span a wide range of natural variation within the microbiome. **(A)** Landscape rarity was measured as the percentage of rosemary scrub patches occupied across the landscape. **(B)** Habitat specialization on the rosemary scrub was measured as the number of rosemary scrub sites occupied divided by the number of rosemary and flatwood sites occupied (ranging from rosemary specialist (Specialization Index=1) to non-specialist (SI=0.5), with a few isolates somewhat specializing on the other habitat type (SI<0.5)). **(C)** Degree centrality – how connected each taxa is within the microbial network – was measured as the number of taxa linked to a microbe in the network divided by the total number of taxa in the network. The schematic (left) is a diagrammatic representation of a microbial co-occurrence network with highly-connected “central microbes” in yellow, “intermediate microbes” in green, and “peripheral microbes” in blue. For all panels, bars represent the percent of microbes in the overall natural rosemary scrub community that fall within each bin (based on our independently-collected NGS dataset across 64 sites). Bold numbers or dashed lines indicate unique isolated taxa (biological replicates) selected as “early colonizers” in the experiment that fall within each bin.

We found that network connectivity is an important indicator of an early colonzier’s role in supporting prokaryotic biodiversity and is a better predictor of diversity and richness than the other ecological attributes of landscape rarity and degree of habitat specialization. By 7 and 14 days post-inoculation, early colonizers with greater centrality had more diverse (F_1,37_ = 11.0, p = 0.002; Fig 2A) and richer (F_1,37_ = 10.6, p = 0.002; Fig 2B) communities. In fact, centrality of early colonizers explained ∼25% of the variance in microbial diversity (rho = 0.48, p = 0.002) and ∼20% of the variance in richness (rho = 0.42, p = 0.008) across all samples (Fig 2 A-B). These results demonstrate that highly-central microbes can act as keystone taxa during early succession by increasing community-wide diversity, which has been linked to greater microbiome stability, resistance to perturbations, and soil functional capacity^12^.

**Fig 2.**
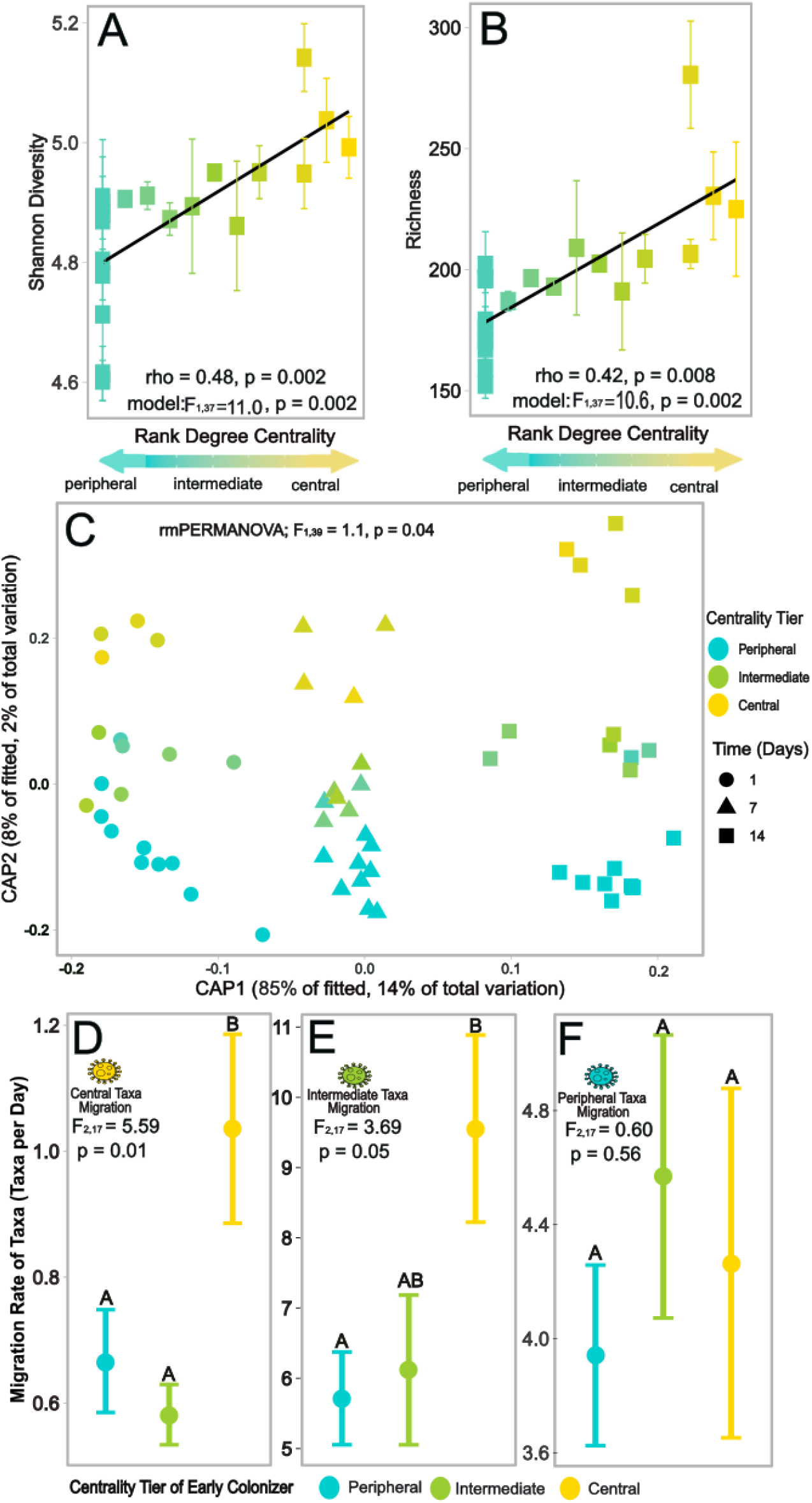
Centrality of early colonizing microbes affected early successional communities’ diversity, richness, composition, and recruitment of other structurally-important, central taxa. **(A, B)** Shannon diversity and richness of assembling soil microbiomes increased with degree centrality of the experimentally inoculated early colonizer. Each point represents one early colonizer used in the manipulative field experiment, and error bars indicate variation among time points (standard error). Statistics are given for Spearman correlations and repeated measures linear models. **(C)** Community composition was impacted by the centrality of early colonizers as can be visualized in this db-RDA ordination of prokaryotic community composition, time, and degree centrality. Each point represents the prokaryotic community composition of a microcosm inoculated with different early colonizers. Points are colored by early colonizer centrality and shapes represent the collection time (days post inoculation). Statistics are given for the repeated-measures PERMANOVA using degree centrality of early colonizers as the explanatory variable. **(D-F)** Centrality of early colonizers also affected the network structure of the microbiome with central early colonizers recruiting significantly more central and intermediate microbes to their communities while peripheral recruitment did not differ. The migration rates (number of taxa in assembled community divided by total days) of the assembling central **(D)**, intermediate **(E)**, and peripheral **(F)** taxa are depicted across the microbiomes for each centrality tier of the early colonizer. Points are colored by centrality tier, error bars indicate standard error, and different letters denote significant differences (p<0.05) from an ANOVA followed by a posthoc Tukey test.

These keystone effects also extended to microbiome composition where differences in degree centrality of early colonizers impacted community-wide structure (repeated-measures PERMANOVA; F_1,39_ = 1.1, p = 0.04; Fig 2C). Despite replicates within each centrality tier being biologically unique (i.e. different early colonizer taxa), the assembled communities clustered by the colonizer’s degree centrality. Importantly, this indicates that central microbes predictably create similar communities regardless of their taxonomic identity. While patterns of succession observed across diverse microbial communities have often been attributed to habitat filtering^4^, our results suggest that connectivity of the early colonizer also plays an important role in the predictability and trajectory of succession.

In addition to enhancing diversity and restructuring community makeup, central early colonizers likely inhibited stochasticity during community assembly by increasing recruitment of other central microbes. We compared how central, intermediate, and peripheral early colonizers impacted recruitment of other microbes in these categories during early succession (see details in Supplementary methods). Interestingly, central early colonizers recruited 66% more central taxa to their newly assembling communities per day than less-connected early colonizers (ANOVA; F_2,17_ = 5.6, p = 0.01) (Fig. 2D), indicating that these central colonizers structure communities by actively recruiting other potentially influential microbes. These effects on the non-random recruitment of other taxa is further demonstrated by the marginally significant increase in migration of intermediate microbes into microcosms with central early colonizers compared to intermediate and peripheral early colonizers (ANOVA; F_2,17_ = 3.7, p = 0.05) (Fig. 2E). In contrast, the migration of peripheral taxa was independent of early colonizer centrality (ANOVA; F_2,17_ = 0.6, p = 0.56)(Fig. 2F), validating that peripheral taxa are acting as transient microbes minimally affected by other microbes. Taken together, these results demonstrate that central early colonizers deterministically impact early community assembly by increasing biodiversity through active recruitment of other influential, central taxa rather than stochastically recruiting a broader array of random, transient microbes.

In conclusion, we combine culturing, sequencing, network theory, and manipulative field experiments to demonstrate for the first time that centrality is a powerful indicator of which microbial taxa are keystone species in nature. Importantly, central early colonizers: 1) increased biodiversity, 2) structured communities, and 3) deterministically recruited a more connected community during early succession. Our work demonstrates that the central placement of hub taxa within community networks can be used to successfully identify structurally significant microbes in complex communities and opens new opportunities for effectively engineering microbiomes through intermicrobial interactions.

## Data Availability

Data is available at Zenodo and will be open access upon publication.

## Acknowledgements

We thank the C. Searcy lab, L. Müller, and W. Browne for feedback on the paper, the UMGC for sequencing support, and E. Fusco, A. Parra, and E. Parra for laboratory assistance. We also thank

A. David, E. Menges, and Archbold Biological Station for enabling us to conduct our field work in the imperiled Florida Rosemary Scrub. Research was funded by the University of Miami research funds and the National Science Foundation (DEB-1922521, DEB-2030060) awarded to M. Afkhami. A. Rawstern was supported by the University of Miami Department of Biology and the Lisa D. Anness Graduate Fellowship, and D. Hernandez was supported by the Maytag Fellowship from the University of Miami.

