## Supplemental Methods for "Hub taxa are keystone microbes during early succession"

### *Study system*

This study was conducted in the Florida Rosemary Scrub ecosystem at Archbold Biological Station on the Lake Wales Ridge (Venus, FL, USA). This ecosystem has a high rate of endemism and supports many imperiled endemic plant and animal species[1]. Florida Rosemary Scrub is characterized by open sand gaps between the dominant allelopathic Florida rosemary shrub (*Ceratiola ericoides)* and occurs as isolated patches surrounded by flatwoods. The soil microbiome of the rosemary scrub is distinct from the surrounding flatwoods habitat[2] and is important for the persistence of imperiled plant populations[3]. The Florida Rosemary scrub is also a pyrogenic ecosystem with natural and prescribed fires that disrupt microbial soil communities by sterilizing the top layer of soil[4]. Recent work has shown that the pulse disturbance effects of fire can alter microbial community composition with cascading effects on germination rates of native plants at early colonization stages[5]. Due to the importance of this habitat, its distinct microbiome, and the disturbance effects of fire, this is an excellent system for studying how attributes of early microbial colonizers affect microbial community succession.

### *Isolation and identification of rosemary scrub microbial taxa*

We isolated microbial taxa from the Florida Rosemary Scrub using soils we collected across Archbold Biological Station in July 2017[2]. We suspended 1 gram of soil in 50 mL of sterile water, serially diluted to a final concentration of $10^{-5}$ µg/µL, and plated 200 µL of solution onto 48 yeast extract agar plates. To increase diversity of our isolated taxa, we varied the growth conditions in a factorial design with 4 carbon sources (mannitol, glucose, sucrose, maltose), 3 pH levels (6, 7, 8), and 4 incubation temperatures (23°C, 28°C, 37°C, 55°C). Sugars were filter-sterilized at a 10% concentration and added after the media was autoclaved to prevent oxidation of sugars. Isolated taxa were cataloged for unique morphological characteristics and purified in liquid media with the same growth conditions. Isolated taxa were preserved as glycerol stocks[6] (20% glycerol v/v) and stored at -80°C. In total, we generated 66 morphologically unique isolated taxa.

To determine the identity of each of the 66 isolated taxa, we extracted DNA for Sanger sequencing. Each of the cultures was grown in 10 mL of liquid media and centrifuged. Washed pellets were transferred to an E.Z.N.A. disruptor tube with SLX-MIus Buffer (OMEGA Bio-Tek, Norcross, GA, USA). Cells were lysed using a TissueLyser II (Qiagen, Carlsbad, CA, USA) at a frequency of 30 $s^{-1}$ for 5 minutes, a heat incubation at 95°C for 10 minutes, followed by 5 additional minutes of bead beating. The DNA was then extracted using the OMEGA Bio-Tek E.Z.N.A. Soil DNA Kit following the manufacturer's protocol. Four negative controls were also included, in which 500 μL of sterile water was used in place of a microbial pellet to test for contamination during the extraction process. DNA quantity was checked using a Qubit 4 fluorometer (Qiagen, Carlsbad, CA, USA) and no cross-contamination occurred with the negative controls. Ribosomal DNA was targeted using primer pairs 515F/806R and ITS7/ITS4 for PCR. We purified the genomic DNA using gel electrophoresis followed by gel extraction (OMEGA Bio-Tek E.Z.N.A Gel Extraction Kit), and 15 μL of purified DNA was mixed with 2 μL 515F (10x dilution) for prokaryotic samples or 2 μL ITS7 (10x dilution) for fungal samples. DNA for each of the 66 samples were Sanger-sequenced at Eurofin Genomics (Louisville, KY, USA).

***Characterization of rosemary scrub taxa's ecological attributes***

We then characterized the role of the isolated microbe within the broader microbial community of this ecosystem using whole community microbiome data from the soil crusts of 103 sites that included 64 rosemary scrub patches and 39 adjacent flatwoods habitats. Soil collection and microbiome sequencing information has been described in detail previously[2]. In brief, samples were collected at each site from the biological soil crust at three sampling points using a sterile soil corer. Those soils were flash frozen in 50 mL conical tubes and stored at −80°C until we extracted the microbial DNA. Total genomic DNA was extracted from 1 g of soil using the OMEGA Bio-Tek E.Z.N.A. Soil DNA Kit following manufacturer's instructions. Fungal (ITS) and prokaryotic (16S) libraries were prepared and sequenced at the University of Minnesota Genomics Center (UMGC) following a dual-indexing approach[7] using Illumina MiSeq (v3, 300 bp paired end, Illumina San Diego, CA, USA). Resulting reads were demultiplexed, merged, and processed through the *QIIME2* pipeline[8] (v2018.8) to remove sequencing adapters, join paired-end reads, and denoise reads. We used *de novo* operational taxonomic unit (OTU) clustering based on 97% sequence similarity[9]. We constructed a rosemary scrub cross-domain microbiome co-occurrence network[10] with *FastSpar*[11] (v1.0.0) with OTUs present in ≥ 10% of all sites and default parameters[12]. *FastSpar*[11] accounts for sparsity in compositional data sets and is a fast and parallelizable implementation of the *SparCC*[13] algorithm with unbiased P-value estimates.

We matched our isolated taxa to the natural microbial community by comparing the Sanger sequences of the cultures to the NGS-generated sequences from across the rosemary scrub community using *BLAST*[14]. Of the 66 isolated taxa sequenced, 20 unique taxa had high quality sequencing data and reliably matched to the microbiome-wide data using the standard e-value criteria[14] (e-value $< 1 \times 10^{-50}$) that we ultimately used as "early colonizers" in our field experiment (described below). To distinguish between top *BLAST* hits with similar quality (i.e. e-values), we chose the top hit found in at least 10% of the sampled field sites as the isolated taxon's identity. If none of the top three *BLAST* hits met this criteria, the microbe identity was assigned as the top *BLAST* hit and the taxon was classified as a transient microbe.

Using this information, we characterized our isolated taxa in three ways: their landscape rarity in the ecosystem (i.e., rosemary scrub occupancy), their degree of habitat specialization, and their centrality within the microbiome network. First, we determined the landscape rarity of each microbe by calculating the proportion of all sequenced Florida rosemary scrub patches (64 patches) that the microbe actually occurs in.

$$landscape\ rarity = \frac{No. of\ rosemary\ sites\ microbe\ occupies}{No. of\ total\ rosemary\ sites}$$

Second, we determined the degree of habitat specialization for each cultured microbe by calculating the relative frequency with which the species occurred in the rosemary scrub habitat compared to the neighboring flatwoods habitat[15]. This habitat specialization index can range from values of 0 (flatwoods specialist) to 1 (rosemary scrub specialist) where 0.5 is a non-specialist and is calculated as:

$$specialization = \frac{No. of\ rosemary\ sites\ occupied}{No. of\ rosemary\ AND\ flatwoods\ sites\ occupied}$$

Third, we assessed the degree centrality of each cultured microbe in the broader microbiome network of the Florida rosemary scrub. Degree centrality measures how connected a node (here microbial taxa) is to the other nodes in the network[16]. Microbial taxa with high degree centrality are the most connected taxa and thus are often considered putative keystones that would influence microbiome structure[17]. Computational removal of these highly connected "hub taxa" has been shown to impact community richness *in silico*[18] reinforcing the prediction that highly-central microbes are keystones. Degree centrality is the number of connections a microbe has within the network divided by how many other microbes are in the network (network size minus one)[19].

$$degree\ centrality\ = \frac{No. of\ taxa\ linked\ to\ target\ in\ network}{No. of\ taxa\ in\ the\ network\ -\ 1}$$

Following previously established methods[20], taxa within the network were binned into centrality tiers by ranking all the microbial nodes in the whole community network from highest to lowest, then partitioning the ranks into central and intermediate microbes. The top 25% of nodes were classified as highly central (predicted keystones) and the remaining nodes were classified as intermediate. OTUs found in ≤ 10% of sites were considered peripheral (i.e. transient) microbes since they did not meet the threshold for network inclusion[12] and were assigned a centrality of 0 since these taxa were not connected to other taxa in the network.

### *Field experiment and microbiome assembly*

To assess how characteristics of early colonizing microbes can influence succession, we set up a manipulative field experiment in which we monitored early microbial community assembly in soil microcosms inoculated with each of our 20 isolated taxa (as well as uninoculated microcosms) at Archbold Biological Station. To create experimental microcosms, sterile pots were filled with 50 g of sterilized sandy soil from the rosemary scrub. This soil was collected from the biological soil crust of the Florida rosemary scrub at Archbold Biological Station (same site as the field experiment) and sterilized at 121°C three times prior to use with at least 24 hours between each sterilization. Under aseptic conditions, microcosms were inoculated with 5 mL of microbial inoculum (1 x $10^6$ cells per mL) from one of 20 early colonizers (i.e., inoculated with one of the 20 isolated taxa suspended in sterile water) or 5 mL of sterile water as a control (21 treatments x 5 replicate microcosms = 105 total microcosms). The microcosms inoculated with one of the 20 isolated microbes ultimately contained 1 x $10^5$ cells per g of soil, which is approximately 1% of the microbial abundance in typical Florida scrub soils as determined by qPCR of rosemary crust soil samples in our broader community sequencing across 64 rosemary scrub habitat patches[2]. This inoculum amount also represents biologically relevant levels of abundance for a single early colonizer species in a natural habitat after a severe fire[21]. All pots received an additional 15 mL of sterile water to distribute the inoculum throughout the soil and prevent desiccation during experiment set up.

Inoculated microcosm pots were wrapped in sterile foil and immediately transported to Archbold Biological Station rosemary habitat patch '1N' (Latitude 27.20, Longitude -81.36). Microcosms were deployed in a completely randomized design within an open sand patch at least 0.5 m away from Florida rosemary shrubs (*Ceratiola ericoides*) to avoid effects of allelopathy[22]. The microcosms were buried 1 cm into the field soil (i.e., the depth at which fire can sterilize soil in this habitat) so that the openings on the bottoms of microcosms were in contact with the rosemary crust for potential microbial migration from below as well as from the air[4]. The openings at the bottom of the microcosms also prevented them from becoming unnaturally water-logged when it rains. Three hundred and fifteen soil samples were collected in sterile 2 mL tubes across 3 time points during early succession (1 day, 7 days, and 14 days after being placed in the field). All soil samples were stored at -80°C until we extracted the DNA of the soil microbiomes.

***Microbiome DNA extraction and sequencing***

DNA was extracted from each soil sample ($n$ = 315; 21 treatment groups, 3 time points, 5 replicates) using the DNeasy PowerSoil Pro QIAcube HT Kit following protocols described in detail elsewhere[5]. Briefly, for DNA extraction of each sample we used 550 mg of homogenized soil and 550 µL of sterile water for negative controls. DNA was quantified with a Qubit 4 fluorometer and concentrations were normalized to 5 ng/µL. All negative controls had undetected fluorometer readings indicating contaminants were not present during the extraction process. Prokaryotic (archaea/bacteria) DNA was targeted using primer pairs 515F/806R for PCR. Libraries were prepared for sequencing using a two-step dual indexing protocol[7]. After each PCR step, magnetic bead cleaning was performed and DNA quality was checked using 1% agarose gel electrophoresis. Indexed DNA from all 315 field experiment samples were pooled in equimolar quantities. Libraries were sequenced on an Illumina MiSeq Sequencer (v3, 300 bp paired end) at the University of Miami Center for Genome Technology (Miami, FL, USA). Sequencing primers[5] were used that matched the universal tail sequences from the first round of amplification.

Sequences were processed through *QIIME2* (v.2022.2) to join paired-end reads, remove low quality bases, and classify OTUs into "species"[8]. Denoising was performed with the *DADA2* algorithm[23], which removes chimeric sequences and truncates amplicon forward and reverse reads to an equal length. Then open reference clustering with the *SILVA* v.132 database was used to group OTUs into species based on 97% sequencing similarity. All OTU abundance tables from *QIIME2* were imported into R v4.1.1[24]. Replicates were pooled and rarefied (to 6000 reads) based on rarefaction curve analysis (R package *GUniFrac*[26], v1.5). Following best practices[25], sequences with relative abundances of less than 0.25% were removed to avoid overinflating diversity due to spurious sequences.

***Microbial diversity, community composition, and community connectivity analyses***

Prokaryotic Shannon diversity, richness, and evenness were calculated in the R package *vegan*[27] (v2.6-2). To determine which early colonizer attribute(s) affected community diversity, model selection was employed using the *dredge* function (R package *MuMIn*[28] v1.44.3). The fixed effects terms considered in the full model were landscape rarity, specialization index, and degree centrality of the early colonizing microbe the microcosm received, as well as the random effects of time point of microbiome collection and all interactions with time. The model with centrality of the early colonizer as the only predictor was the best model (lowest delta AICc; Figure S1)[29]. The estimated proportion of variance explained by the fixed effects in the model was calculated using a Spearman correlation.

After finding that centrality was the most important attribute predicting prokaryotic diversity and richness, we conducted follow-up analyses on how this attribute affected composition of the community. To do this, we conducted a distance-based redundancy analysis (db-RDA) using Aitchison dissimilarity distances[30], which is recommended for microbiome data[31]. A pseudo-count of one was applied when using the centered log ratio (clr) transformation for the Aitchison distance. The terms used in the db-RDA were degree centrality and time. Repeated measures PERMANOVA was used to assess whether the degree centrality gradient was significant across all time points. The db-RDA analysis and rmPERMANOVA were performed with the *capscale* and *adonis2* functions, respectively (R package *vegan*[27] v.2.6-2).

To understand how early colonizer centrality impacted migration of other microbes into the assembling communities, all the OTUs from the assembling microbiomes within our experimental microcosms were matched using *BLAST* to the OTUs from our rosemary scrub microbiome network (based on the 64 field sites). This allowed us to extract the degree centrality of each microbe that migrated into the microcosms. Taxa from the assembled communities were then binned into centrality tiers using the same method employed earlier (described in the "characterization of rosemary scrub taxa's ecological attributes" section). Briefly, the degree centrality values from the sequenced microcosms were ranked from highest to lowest and partitioned such that taxa with the top 25% of degree centralities were classified as central, the taxa with non-zero degree centralities were classified as intermediate, and the taxa with zero degree centralities were classified as peripheral. The migration rates for central, intermediate, and peripheral microbes into the assembling communities were each calculated by dividing the number of taxa observed in the assembled communities of that centrality tier by the total number of days (14 days) to estimate the average recruitment of new taxa to the communities per day for each centrality tier. ANOVAs with *post hoc* Tukey's HSD tests were performed to compare the migration rates of the microbes between microcosms inoculated with early colonizers with different centrality tier attributes.

**Supplementary Figures:**

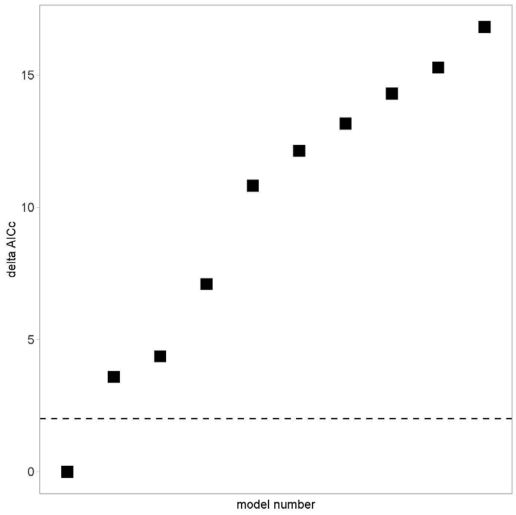


**Supplementary Fig 1: Comparison of the top ten best models from global model selection.** Only the best model (Shannon diversity ~ centrality of early colonizer) had a delta AICc under the threshold of 2.
